# Long-term monitoring reveals topographical features and vegetation explain winter habitat use of an Arctic rodent

**DOI:** 10.1101/2021.01.24.427984

**Authors:** Xaver von Beckerath, Gita Benadi, Olivier Gilg, Benoît Sittler, Glenn Yannic, Alexandra-Maria Klein, Bernhard Eitzinger

## Abstract

Collapsing lemming cycles have been observed across the Arctic, presumably due to global warming creating less favorable winter conditions. The quality of wintering habitats, such as depth of snow cover, plays a key role in sustaining population dynamics of arctic lemmings. However, few studies so far investigated habitat use during the arctic winter. Here, we used a unique long-term time series to test whether lemmings are associated with topographical and vegetational habitat features for their winter refugi. We examined yearly numbers and distribution of 22,769 winter nests of the collared lemming *Dicrostonyx groenlandicus* from an ongoing long-term research on Traill Island, Northeast Greenland, collected between 1989 and 2019, and correlated this information with data on dominant vegetation types, elevation and slope. We specifically asked if lemming nests were more frequent at sites with preferred food plants such as *Dryas octopetala x integrifolia* and at sites with increased snow cover. We found that the number of lemming nests was highest in areas with a high proportion of *Dryas* heath, but also correlated with other vegetation types which suggest some flexibility in resource use of wintering lemmings. Conversely, they showed a distinct preference for sloped terrain, probably as it enhances the formation of deep snow drifts which increases the insulative characteristics of the snowpack and protection from predators. With global warming, prime lemming winter habitats may become scarce through alteration of snow physical properties, potentially resulting in negative consequence for the whole community of terrestrial vertebrates.

## Introduction

Climate change is impacting the Earth’s ecological and biological systems. According to the Intergovernmental Panel on Climate Change (IPCC), the Arctic surface temperature has increased twice as fast as the global average over the last 20 years, with alterations in precipitation, snow cover and permafrost temperature (Pörtner et al. 2019). Environmental change has triggered measurable shifts in performance and distributions of plant and animal species, translating into ecosystem-wide changes (Post et al. 2009). Arctic ecosystems are hypothesized to have little functional redundancy (Wookey et al. 2009) and changes in phenology, distribution and behavior of species may therefore have disproportional impacts on the structure and functioning of their communities and ecosystems (Gilg et al. 2012). Among vertebrates, lemmings play a key role in the High Arctic tundra food web (Batzli et al. 1980). They are year-round residents and thus have to survive on local primary production throughout the year (Soininen et al. 2015). In addition, they show little migratory behavior which emphasizes the importance of local vegetation for their survival (Gilg 2002). As a key prey for many terrestrial arctic predators, lemmings impact reproduction and abundance of those, which in turn creates feedbacks on lemming cyclic dynamics (Sittler et al. 2000, Gilg et al. 2003, 2006, Therrien et al. 2014). Lemmings also influence the dynamics of other vertebrate prey such as geese (Bêty et al. 2001) and other incidental prey species (Mckinnon et al. 2013). Lemmings are well-known for their regular, large-amplitude population cycles (Ims and Fuglei 2005), climaxing every three to five years (Gilg et al. 2003, Gruyer et al. 2008, Ehrich et al. 2020). Although predator-prey interactions are commonly argued to be the cause of the lemming cycles (Gilg et al. 2003, Fauteux et al. 2016), the role of the environment in the modulation of such dynamics is only poorly understood (Kausrud et al. 2008). For example, population cycles of the Norwegian lemming *Lemmus lemmus* have been found to be driven by the availability of food plants (Turchin et al. 2000), however, recent studies have challenged this assumption (Ims et al. 2011, Soininen et al. 2015).

Collapsing lemming cycles have been observed at different sites in the Arctic, including Northeast Greenland (Kausrud et al. 2008, Gilg et al. 2009, Schmidt et al. 2012b, Ehrich et al. 2020). The most prevailing explanation for these disruptions is the impact of changing snow characteristics due to climate change (Ims et al. 2008, Kausrud et al. 2008, Gilg et al. 2009). Thick snow cover insulates wintering lemmings from low ambient air temperatures and increases their protection from predators (Reid et al. 2012). Thus, a thin snowpack appears to be a strong limiting factor for recruitment and lemming survival, that may maintain lemming at low densities (Reid and Krebs 1996). Moreover, habitat selection studies suggest that climate-induced landscape change may be beneficial for some lemming species and detrimental for others (Morris et al. 2011). The detrimental effect of global warming is supported by a paleogenomics study that shows previous climate warming events to have strongly reduced the population size of arctic lemmings *Dicrostonyx torquatus* (Prost et al. 2010). Furthermore, according to ecological niche modeling, arctic lemmings lost approximately 80% of their suitable habitat since the last glacial period, 21,0000 years ago (Prost et al. 2013).

The collared lemming *Dicrostonyx groenlandicus* is found in High Arctic ecosystems of northern Canada, Alaska and Greenland. Dry dwarf-shrub heath and herb slopes, which are rich in *Salix arctica* and *Dryas spp*., are the preferred summer habitats of the collared lemming (Batzli et al. 1983, Schmidt 2000, Predavec and Krebs 2000, Morris et al. 2011, Ale et al. 2011). However, habitat use during winter remains poorly known due to the challenges involved in studying them beneath the snow in the harsh winter arctic conditions (Soininen et al. 2015). Yet, successful reproduction during the winter season plays a key role in lemming population dynamics (Ims et al. 2011, Bilodeau et al. 2013a) and it is even considered a necessary condition for the occurrence of a population outbreak (Fauteux et al. 2015). Lemmings build subnivean nests (i.e. nests under the snow) made of vegetation, which are assumed to provide insulation against low temperatures and to protect litters from predation (Sittler 1995, Duchesne et al. 2011, Domine et al. 2018).

Winter nests are generally built on the ground or within the snowpack. After snowmelt, these nests can be found lying on the ground with no signs of connection to the vegetation below. Winter nests are often clumped and connected through tunnels, suggesting some social relationship between their inhabitants (Sittler 1995). Collared lemmings stay in their winter quarters until the snowpack is soaked with melt water and collapses. By that time, lemmings seek their summer habitats in dry and early snow free heath lands (Fauteux et al. 2018). So far, surprisingly few studies have investigated these nests to identify habitat usage during winter, even though winter nests are easily spotted after snowmelt in the Arctic tundra.

To date, the most comprehensive study exploring the effects of snow cover and habitat features on the distributions and reproduction of winter nests took place on Bylot Island, Canada (Duchesne et al. 2011). Here, collared lemming nests, sampled over two consecutive years, were mainly located in areas with high micro-topography heterogeneity, steep slopes, deep snow and a high cover of mosses (Duchesne et al. 2011). Other studies focused solely on the relationship between snow cover and winter nests. These studies noted a concentration of winter nests in areas of greatest winter snow accumulation (Reid and Krebs 1996, Reid et al. 2012, Bilodeau et al. 2013a), in particular on sloped terrain, which provides shelter for the deposition of wind-blown snow in drifts (Reid et al. 2012). An alternative approach to analyzing winter habitat preferences of lemmings is to identify plant fragments in their stomach contents or faecal pellets. Such studies documented that collared lemmings feed mostly on dicots during the arctic winter (Batzli et al. 1983, Rodgers and Lewis 1985, Bergman and Krebs 1993), specifically *Dryas spp*. and *S. arctica* (Berg 2003). Winter diets of lemmings were dominated by different species of *Salix* in a more recent study based on environmental DNA metabarcoding of faecal pellets, and mosses, which are the most dominant plants in locations with many lemming nests, are of minor importance (Soininen et al. 2015). However, several older studies found that the lemming’s diet varied between sites, which suggests some flexibility in food choice (Rodgers and Lewis 1985, Bergman and Krebs 1993).

Collectively, these studies demonstrate that no clear pattern has been detected to explain how lemmings choose their winter habitat. This could be due to the fact that past studies exclusively captured individual population phases (Duchesne et al. 2011, Reid et al. 2012, Soininen et al. 2015), possibly ignoring the effect of a cycling population on habitat use, i.e. a potential density-dependent effect. The assumption is that lemmings undergo density-dependent dispersal as a result of intraspecific competition in the subnivean space and may be forced to emigrate to less optimal habitats, as preferred sites are thought to be limited. This study is the first to examine the long-term determinants of winter habitat use of lemmings by analyzing data collected over 30 years. Using this unique time series, we were able to define characteristics of lemming winter habitats at an unprecedented scale. The objective of our study is to determine the ecological factors influencing the spatial distribution of winter nests of collared lemmings, allowing to identify areas of frequent habitat use. Habitat use is defined as the way an animal consumes a collection of physical and biological components in a habitat, while habitat selection takes into account the availability of resources (Hall et al. 1997), which we were not able to consider in our analysis. We hypothesize that lemmings mainly use areas providing a deep snow cover and a rich plant food source. As *Dryas* heath hosts the highest abundance of preferred food plants, we predict that, (1) sites with a greater abundance of *Dryas* heath and greater vegetation cover have a positive effect on nest counts. Furthermore, since snow depth is highest at the bottom of steep slopes, we predict that (2) the number of lemming winter nests is higher on sites with increased slopes.

## Material and Methods

### Study area

This study was carried out as part of the joint French-German research project “Karupelv Valley Project” (https://www.karupelv-valley-project.de/english; http://www.grearctique.org/karupelv). The core research area covers approximately 15 km^2^ and is located on the southwestern coast of Traill Island, Northeast-Greenland (72.5°N; 24°W, Fig. 1). The study area is bordered by the Karupelv river to the south and the Eskdal river to the east (Fig. 1). The northern border follows small streams and geomorphological structures towards the coast (Rau 1995). The region is part of the Northeast Greenland National Park and offers a study site with minimal human impact for long-term surveys (Sittler 1995). Mean summer temperature in July is 5.8°C and mean winter temperature in February −22.4°C. Annual precipitation is 261 mm with relatively dry summers as most falls as snow in winter (Hansen et al. 2008). Snow cover is usually present from September to June or July (Gilg et al. 2009). The study area covers lowlands and sloped terrain of different scales, the maximum slope gradient is 67° and the area spans elevation ranges from 0 to 122 m a.s.l. (Porter et al. 2018).

**Figure 1.**
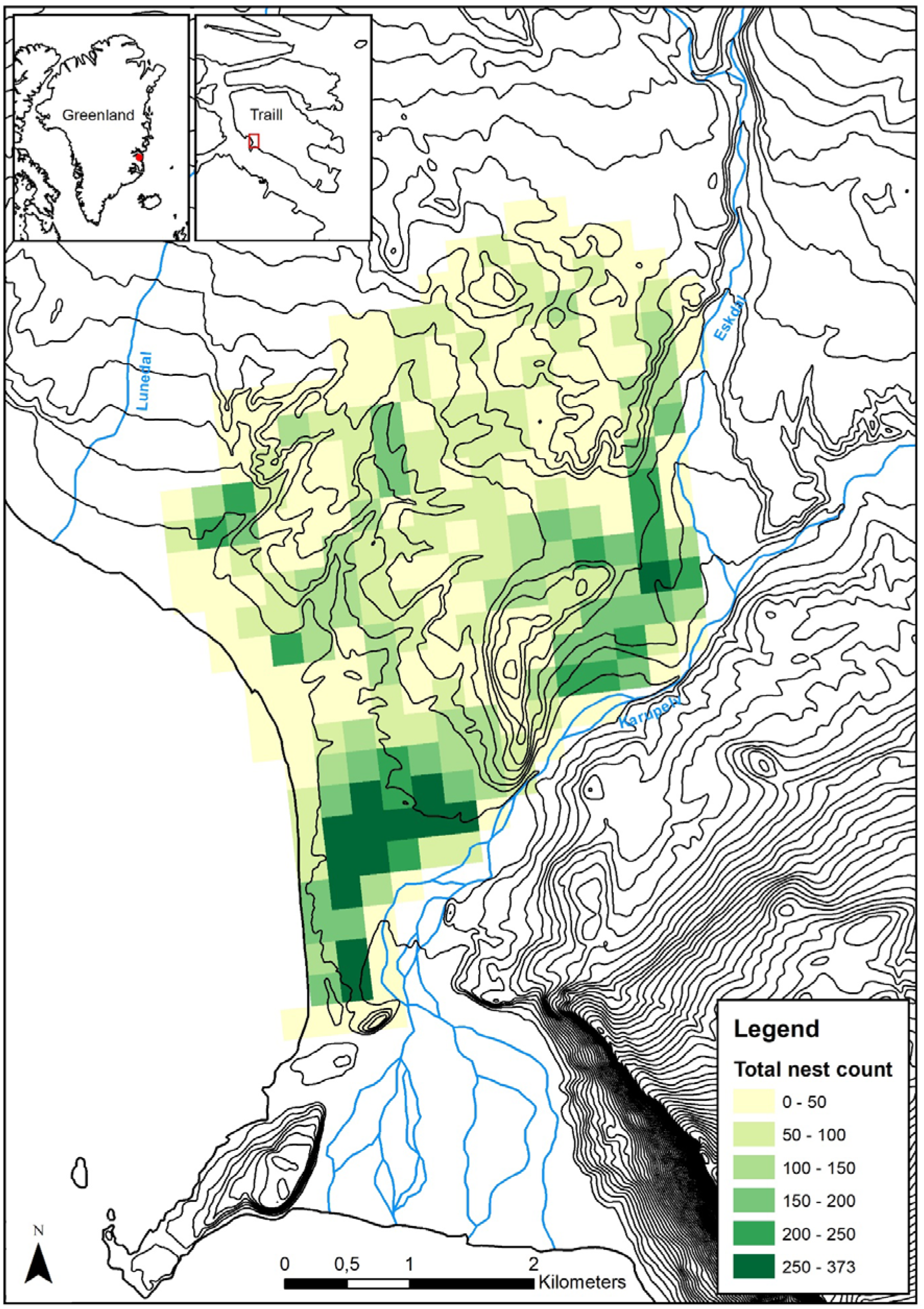
Cumulated number of lemming winter nest censused between 1989 and 2019 on 274 grids of ca. 250×250 m in the Karupelv valley on Traill Island, Northeast-Greenland.

The site is classified as High Arctic tundra (Walker et al. 2005), and has five dominating vegetation types (Rau 1995). *Cassiope* Heath consists of pure stands of *Cassiope tetragona* and *Dryas* Heath is dominated by *Dryas octopetala x integrifolia*. Spot tundra is a sparse vegetation with sporadically occurring vegetation islands of *Cassiope* heath and *Dryas* heath. In areas with wet conditions during the whole vegetation period such as streams and fens, a tundra-typical wet vegetation occurs, dominated by mosses and relatively tall sedges (*Carex* spp., *Eriphorum scheuchzeri*). Characteristics of the recent floodplains are various species of rockfoils (e.g. *Saxifraga aizoides*). The vegetation-free areas resemble the semi-polar deserts in the North and are characterized by sand dunes (Rau 1995). The dominant mammal species is the collared lemming, which forms the main prey for arctic fox *Vulpes lagopus*, stoat *Mustela erminea*, snowy owl *Bubo scandiacus* and, during peak years, also for long-tailed skua *Stercorarius longicaudus* (Gilg et al. 2006, Andreassen et al. 2017).

### Winter nest survey

To document lemming cycles in a long-term perspective, the research project opted for a yearly monitoring based on the non-invasive and easily replicable census of winter nests. These nests are obligate indicators for the prolonged stay of the lemmings in the subnivean environment and can therefore provide information on their habitat use in the Arctic Tundra.

Although several lemmings can inhabit the same nest (Rodgers and Lewis 1986), winter nests are a good proxy of lemming population densities (Gilg et al. 2006). Winter nests were recorded by the same person (BS) from 1988 to 2019. The census took place between the end of June and the beginning of August, i.e. for about four weeks after snowmelt. Winter nests are easily detectable after snow melt in the High Arctic tundra and can thus be counted accurately over a larger area (Sittler 1995). Since nests need to be destroyed upon recording to avoid double counting in the following years, the first year of data assessment was omitted. Hence, this study includes 31 years of data (1989-2019). Nests were recorded while walking 30 m-wide transects on flat terrain, and 20 m-wide in rugged topography. High quality aerial photography was used to divide the study region into grid cells and ensure that the same census effort was devoted to each cell (Sittler 1995). The same aerial photography was then used to assign nest records to a specific cell. For that, the study region was divided into squares of ca. 250 × 250 m. This methodology gives indices on relative abundance, habitat use and spacing patterns during winter (Sittler 1995).

### Data processing

Answering our research questions required the revision of the nest data (see Supporting information) and the creation of a digital version of the above-mentioned grid. GIS-based data management and extraction was implemented with ArcMap 10.7.1. First, all aerial photographs were georeferenced with a high-resolution (10 m) satellite image (see Supporting information) from the Copernicus Mission (Copernicus Sentinel data 2017). Next, a new digital grid was created with the fishnet tool that matched the analogue one in a best-fit scenario. As the aerial photographs did not align perfectly, some deviation was expected. Many of the contiguous aerial photographs slightly overlapped with each other (see Supporting information), thus it was necessary to manually merge certain cells and create a new grid identification system (see Supporting information). Nest counts were translated to the new identification system and assigned to the digital grid. Single cells, which were located outside the core study area and did not contain any nest recordings, were erased. Therefore, the new grid consists of 274 cells of 250 × 250 m (62,500 m^2^) in size, with a total extent of approximately 15 km^2^.

Elevation (m a.s.l.) and slope (degrees) were extracted from a digital elevation model (5 m resolution) supplied by the ArcticDem project (Porter et al. 2018). Elevation units are referenced to the WGS84 ellipsoid by the ArcticDem project (Porter et al. 2018). We used the EGM2008 geoid undulation model to convert ellipsoidal height to mean sea level by using the coordinates of the research area (72.5°N; 24°W) as input variables (Pavlis et al. 2012). With the elevation and the digital grid in place we could extract topographic and vegetation data for each patch using common GIS tools. Slope was calculated using the Spatial Analyst toolbar and the digital elevation model. Elevation and slope were then calculated for each patch by taking the mean of all raster cells in that patch (see Supporting information). Vegetation data was derived from a geo-ecological study of the study region (Rau 1995)(Supporting information). This study analyzed vegetation types at the end of the growing season in early August. It classified 13 different vegetation types throughout the core research area. For a more detailed description of the vegetation types see Supporting information. Given the slow changes in vegetation in the region (Schmidt et al. 2012a), we considered that this dataset was relevant to use for the entire study period.

### Statistical analysis

Due to the high inter-annual fluctuations in winter nest numbers, the proportion of empty cells varied greatly between years (see Supporting information for a complete list of nest presence and empty cells). We first fitted a Generalized Linear Model (GLM) with Poisson errors and log link to the data, but since diagnostic tests indicated significant overdispersion and zero-inflation, we used a zero-inflated negative binomial GLM as our final model, with only an intercept fitted to account for the excess number of zeros. Additionally, we accounted for possible non-independence of data points by testing for spatial autocorrelation of model residuals separately for each year and for temporal autocorrelation separately for each grid cell. Since spatial autocorrelation was detected in 23 of 31 years and temporal autocorrelation in 20 of 274 grid cells (see Supporting information), we subsequently applied a Matérn covariance structure to account for spatial autocorrelation and an autoregression structure (AR1) to account for temporal autocorrelation. In the model, the number of lemming winter nests per grid cell was the response variable and mean elevation per cell, mean slope per cell and the area of each vegetation type in the cell were explanatory variables. All variables are continuous. To test for density-effects on habitat use, we also calculated separate models for years with low (< 2 lemmings/ha) and high (> 2 lemmings/ha) lemming densities (Gilg et al. 2019).

Due to the large number of different vegetation types, only the six dominant ones (spot tundra, moss-sedges tundra, *Dryas* heath, *Cassiope* heath and Sand Dunes) were included in this study (see Supporting information). All predictors were standardized to a mean value of 0 and a standard deviation of 1, to facilitate model convergence and enable us to compare the explanatory contributions of each independent variable regardless of units. Testing all predictors for correlation produced correlation coefficients smaller than 0.41. Simulation-based standardized residuals were used to check for patterns in the residuals against fitted values and predictors. All statistical analyses were performed with R 4.0 (R Core Team 2020) and the packages glmmTMB (Brooks et al. 2017) and DHARMa (Hartig 2017).

## Results

In total, 22,769 individual nests were counted on 274 grid cells during 31 years. Seven of these cells (2.5%) did not host any nests at any time, while the highest cumulative number recorded in one cell reached 373 nests. We found no noteworthy difference of habitat use between years of low and high lemming density (Supporting information). Therefore, we here present the results of the model including all years, despite the large difference in total lemming density (Table 1). Six of eight factors in our model had a significant effect on lemming nest numbers. The number of nests was positively related to three vegetation types: Spot tundra (Fig. 2a), moss-sedges tundra (Fig. 2b) and *Dryas* heath (Fig. 2c). Conversely, the number of lemming nests declined with an increase in floodplain vegetation (Fig. 2d), while we found no effect for *Cassiope* heath and sand dunes. Regarding geophysical variables, slope (mean: 4.21 % ± 0.161 SE) increased the number of nests (Fig. 2e) while elevation (mean for all grids: 50 m; range: 0-119 m) had a negative effect (Fig. 2f), meaning that fewer nests were found at high elevation.

**Figure 2:**
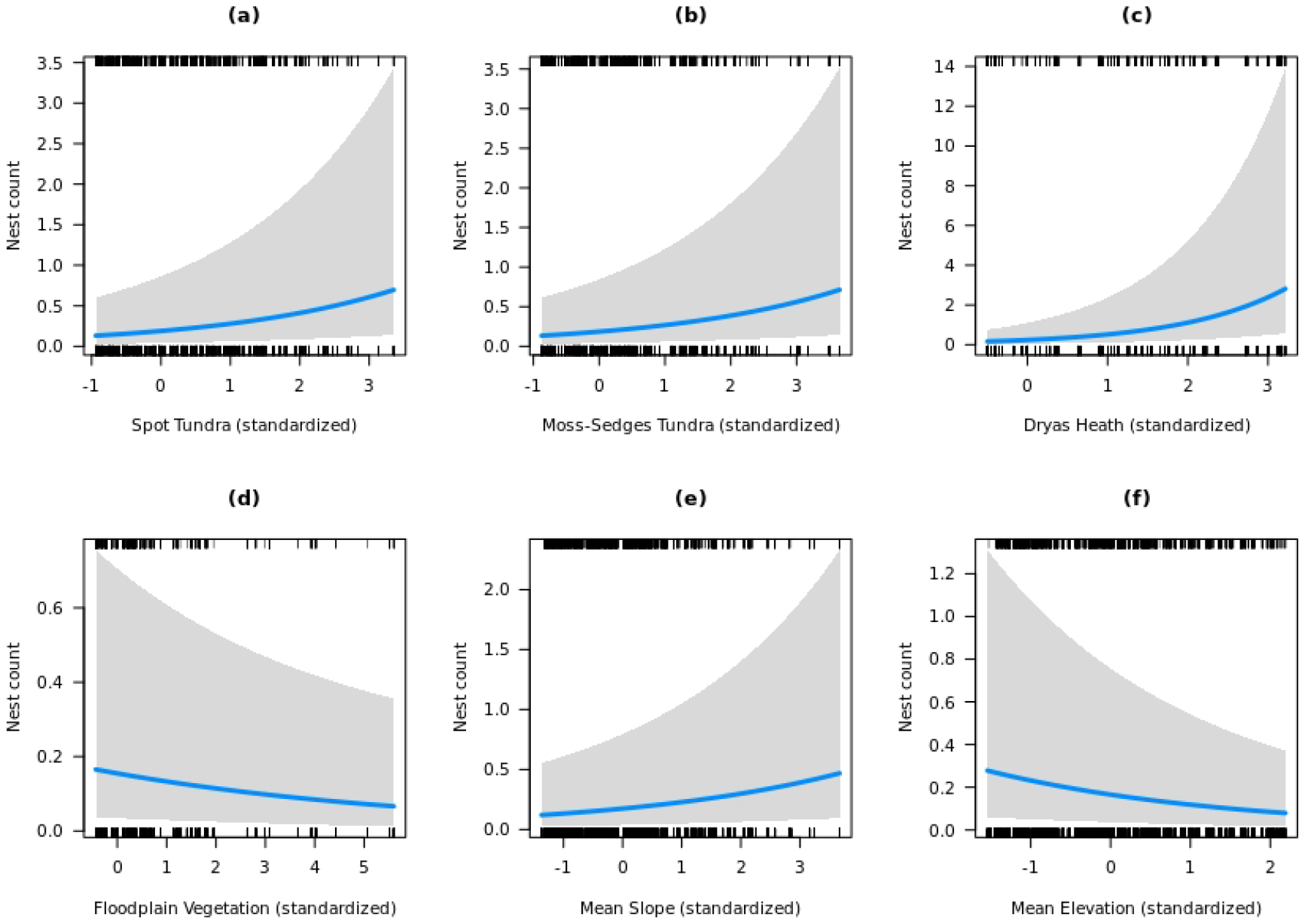
Effect plots for all significant predictors (see Table 1) across all years (1989-2019). Nest counts are shown per grid. Shaded area represents 95% confidence intervals.

**Table 1.** Summary table of negative binomial generalized linear model of the effect of vegetation type, elevation and slope on total lemming nest number for all years (1989-2019). Significant results are highlighted in bold.

| Fixed Factors | Estimate | Std. Error | Z-value | P-value |
| --- | --- | --- | --- | --- |
| Intercept | -0.39752 | 0.22342 | -1.779 | 0.0752 |
| **Spot tundra** | **0.38838** | **0.07608** | **5.105** | **<0.001** |
| **Moss-sedges tundra** | **0.36903** | **0.06975** | **5.291** | **<0.001** |
| **Dryas heath** | **0.76331** | **0.08507** | **8.972** | **<0.001** |
| Cassiope heath | 0.06993 | 0.06196 | 1.129 | 0.2591 |
| Sand dunes | -0.03833 | 0.06954 | -0.551 | 0.5815 |
| **Floodplain vegetation** | **-0.15149** | **0.06853** | **-2.211** | **0.0271** |
| **Mean slope** | **0.26964** | **0.06755** | **3.992** | **<0.001** |
| **Mean elevation** | **-0.33578** | **0.08084** | **-4.154** | **<0.001** |

## Discussion

This study is the first to examine the long-term determinants of winter habitat use of lemmings by analyzing data collected over 30 years. Corroborating results from recent short-term studies (Duchesne et al. 2011, Reid et al. 2012, Soininen et al. 2015), we found that both, vegetation types and geomorphological structures play a significant role in driving habitat use of collared lemmings. In addition, we found no evidence for shifts in habitat use between years of high and low population size, suggesting that lemmings are not constrained by habitat resources. Below, we will examine each finding in turn.

Our study shows that collared lemmings are not restricted to one vegetation type during the winter months, but instead use multiple vegetation types. However, we found indirect support for the hypothesis that lemmings mainly use areas with deep snow cover, characterized by higher topographic heterogeneity. Winter nests were more abundant at locations with a greater proportion of *Dryas* heath than in places that were dominated by spot tundra or moss-sedges tundra, which all host a different plant species composition (Supporting information). We can thus accept the first hypothesis that lemming cluster at sites with *Dryas* heath and key food plant species (e.g. *S. arctica, D. octopetala x integrifolia*) therein. This fits into a series of studies across the Arctic that document a preference of *Dryas* but also some flexibility in the use of food resources by lemmings, as diets vary depending on the dominance of different plant species: *Dryas* was preferred by lemmings at Pearce Point (Northwest Territories, Canada), based on faecal pellets collected in early spring (Bergman and Krebs 1993) and in Igloolik (Nunavut, Canada), based on live-trapping in the snowpack (Rodgers and Lewis 1985). By contrast, in regions with low abundance of *Dryas, Salix* was found to be dominant food plant as indicated by the location of nests, associated latrine sites and faecal pellets in Northern Greenland (Klein and Bay 1994) and percentage of shoots clipped after snowmelt in Alaska, respectively (Batzli et al. 1983). These past findings are consistent with the most recent study on Bylot Island (Nunavut, Canada) analyzing faeces of co-existing brown lemmings and collared lemmings (Soininen et al. 2015). Although the winter diet of *D. groenlandicus* was generally dominated by *Salix*, most plant taxa that were identified as forage were consumed in proportion to their availability.

In our study, *Cassiope* heath cover did not affect winter site use by lemmings, although past research identified *C. tetragona* as non-food plant for lemmings in the winter months that was avoided (Soininen et al. 2015). In fact, lemmings were only observed to eat the flowers of *Cassiope* which are available during summer (Berg 2003). Conversely, we expected *Cassiope* heath to be attractive for lemmings as an indicator for snow-beds (Pedersen et al. 2016). However, as *Cassiope* only covers 4% of the total area, we were unable to detect positive or negative effects on lemming densities.

The most comparable study so far listed areas with a low abundance of lichens and a high abundance of mosses as preferred sites for winter nests (Duchesne et al. 2011). Collared lemmings have also been found to consume a greater proportion of mosses during the winter than during the summer (Rodgers and Lewis 1986), possibly because mosses contain fatty acids keeping body temperature at a high level in cold climates (Prins 1982). Although our study also identified lemming nests on grids with a dominant “moss-sedges tundra” vegetation, they were less strongly favored than *Dryas* heath. We found indication that winter nest site use on Traill Island is mostly linked to *Dryas* heath and to a lesser extent to mosses, sedges and spot tundra. The positive correlation with spot tundra (surface coverage between 31% and 60%) suggests that a low vegetation cover does not affect lemming winter site selection at a certain level. However, habitat types with less than 10% vegetation cover are less likely to be inhabited by lemmings: Floodplain vegetation, which is also characterized by a low biomass and low plant species diversity are being avoided by wintering lemmings, while we also found only low winter use of sand dunes habitat.

Previous studies noted heterogeneity of micro-topography as the most important factor determining both summer habitat (Morris et al. 2000) and winter habitat of lemmings (Duchesne et al. 2011). In line with these studies, slope had a positive effect in our model, and the increased usage of topographic features by wintering lemmings is an indirect effect of habitat configuration in the study region. Indeed, in contrast to the relatively flat plateaus and dunes, the steep slopes of gullies and terraces enhance the formation of deep snow drifts (Bilodeau et al. 2013b). Additionally, the hiemal threshold may be reached faster on wind-protected slopes than in less protected areas after the initial winter storms (Reid et al. 2012). Consequently, as the greatest winter snow accumulation attracts lemmings, this has been connected to a concentration of winter nests (Duchesne et al. 2011, Reid et al. 2012, Bilodeau et al. 2013a).

Deeper snow will provide higher and more stable subnivean temperatures (Duchesne et al. 2011), thus reducing the physiological stress that lemmings undergo during winter (Bilodeau et al. 2013c). A high thermal resistance was found in snow drifts and in willow shrubs, thus explaining their suitability as a habitat for subnivean life (Domine et al. 2016). Additionally, snow offers small mammals some mechanical protection against many predators, although the amount of protection depends on the predator species (Berteaux et al. 2017). For example, snow can effectively protect lemmings from Arctic foxes (Duchesne et al. 2011) but less so from stoats (Bilodeau et al. 2013b), probably because their small size allows them to move under the snow through tunnels created by their small rodent prey (Berteaux et al. 2017). Ground depressions with snow accumulations may however also be disadvantageous as melt–freeze events in spring can form hard refrozen crusts or even ice layers near the ground, thereby destroying the subnivean microhabitat (Berteaux et al. 2017). In addition, rain-on-snow events during autumn can harden the basal snow layer causing the same effect (Domine et al. 2018). Thus, weather conditions during the establishment of the snow-bed largely determine the physical properties of the bottom snow layer (Berteaux et al. 2017). These initial conditions could have a lasting impact on the quality of the subnivean space for the whole winter (Domine et al. 2016). The persistent pattern of snow-depth distribution mainly controlled the spatial distribution of vegetation in a snow modelling experiment in Zackenberg, Northeast-Greenland (Pedersen et al. 2016). Here, *Dryas* Heath occurred at significantly lower snow depth than observed for the remaining four other vegetation types (Pedersen et al. 2016). In addition, a snow fencing experiment on Herschel Island (Canadian Arctic) documented that the winter nest density intensified in areas with an artificially increased snow depth. After removing the fences, snow depth and winter nest density in the treatment area returned to pre-intervention conditions (Reid et al. 2012). The duration of the experiment on Herschel Island (one winter) would not have allowed for a shift in vegetation, emphasizing the dominant role of snow conditions in habitat use of lemmings. Winter nest site counts significantly dropped with higher elevations in our study. Here, lower snow accumulation due to greater exposition to wind and lower vegetation cover most likely forbid the establishment of amenable conditions necessary for survival of lemming in winter.

We found no evidence that nest counts are connected to different habitat features in years with high and low lemming population sizes. Our results however indicate, that by using a wide array of different and abundant vegetation types, lemmings find enough winter nesting sites which are able to support even high lemming populations in peak years. Similarly, the study area provides an abundance of geomorphologic features providing thick snow covers needed for thermal insulation and protection of predators. Lemmings populations, including *Dicrostonyx* on Traill Island, are assumed to be top-down controlled by specialist predators such as the ermine (Sittler 1995, Gilg et al. 2003). However, the ongoing warming of the Arctic and coupled shifts in habitat quantity and quality, along with changes in snow quality, are considered to be the prime factors to explain crash of lemming cycles and reduced overall population numbers (Kausrud et al. 2008, Post et al. 2009, Gilg et al. 2009). While the study region of Northeast Greenland has experienced only moderate shifts in vegetation cover (Schmidt et al. 2012a), recent weather extremes (Schmidt et al. 2019) suggest that alterations in snow regime could strongly impact lemming winter habitat use in the future.

## Conclusion

The results of this study highlight the importance of both, vegetation and geomorphological structures as determinants of the spatial distribution of wintering lemming populations. As warming in the Arctic proceeds, lemmings will be confronted with significant changes in quality and quantity of their (still) abundant winter nesting sites. Shifts in vegetation, rain-on-snow events during autumn and melt-freeze events in spring may alter the quality of these refugia and could decrease the survival of lemmings in winter and be responsible for the recent distortions of population dynamics (Domine et al. 2018), which could potentially affect the whole tundra ecosystem (Gilg et al. 2012). Although lemmings do not decline on a global scale, regional disruptions may be the result of large geographical variability in climate, snow physical properties and community composition across the Arctic (Ehrich et al. 2020). Future studies should aim to further investigate the relation of snow properties and lemming winter habitats. This could e.g. be done by applying high resolution local snow models to look at dynamic (snow) versus rather slowly changing (vegetation) environmental parameters. This would allow us to scrutinize where lemmings would be more likely to persist with climate changes, and finally to assess broader impacts on the structure and functioning of polar ecosystems.

## References

Ale, S. B. et al. 2011. Habitat selection and the scale of ghostly coexistence among Arctic rodents. - Oikos 120:1191–1200.

Andreassen, P. N. S. et al. 2017. Gastrointestinal parasites of two populations of Arctic foxes (Vulpes lagopus) from north-east Greenland. - Polar Res. 36:13.

Batzli, G. et al. 1980. The herbivore-based trophic system. An arctic ecosystem: the coastal tundra at Barrow, Alaska Dowden. - Hutchinson & Ross.

Batzli, G. et al. 1983. Habitat use by lemmings near Barrow, Alaska. - Holarct. Ecol. 6: 255–262.

Berg, T. 2003. The collared lemming (Dicrostonyx groenlandicus) in Greenland: Population dynamics and habitat selection in relation to food quality.

Bergman, C. and Krebs, C. 1993. Diet overlap of collared lemmings and tundra voles at Pearce-Point, Northwest-Territories. - Can. J. Zool.-Rev. Can. Zool. 71: 1703–1709.

Berteaux, D. et al. 2017. Effects of changing permafrost and snow conditions on tundra wildlife: critical places and times. - Arct. Sci. 3: 65–90.

Bêty, J. et al. 2001. Are goose nesting success and lemming cycles linked? Interplay between nest density and predators. - Oikos 93: 388–400.

Bilodeau, F. et al. 2013a. Demographic response of tundra small mammals to a snow fencing experiment. - Oikos 122: 1167–1176.

Bilodeau, F. et al. 2013b. Effect of snow cover on the vulnerability of lemmings to mammalian predators in the Canadian Arctic. - J. Mammal. 94: 813–819.

Bilodeau, F. et al. 2013c. The effect of snow cover on lemming population cycles in the Canadian High Arctic. - Oecologia 172:1007–1016.

Brooks, M. E. et al. 2017. glmmTMB Balances Speed and Flexibility Among Packages for Zero-inflated Generalized Linear Mixed Modeling. - R J. 9: 378–400.

Copernicus Sentinel data 2017. For Sentinel data. Tile: S2A_tile_20170731_26XNF_0.

Domine, F. et al. 2016. Seasonal evolution of the effective thermal conductivity of the snow and the soil in high Arctic herb tundra at Bylot Island, Canada. - Cryosphere 10: 2573–2588.

Domine, F. et al. 2018. Snow physical properties may be a significant determinant of lemming population dynamics in the high Arctic. - Arct. Sci. 4: 813–826.

Duchesne, D. et al. 2011. Habitat selection, reproduction and predation of wintering lemmings in the Arctic. - Oecologia 167: 967–980.

Ehrich, D. et al. 2020. Documenting lemming population change in the Arctic: Can we detect trends? - Ambio 49: 786–800.

Fauteux, D. et al. 2015. Seasonal demography of a cyclic lemming population in the Canadian Arctic. - J. Anim. Ecol. 84: 1412–1422.

Fauteux, D. et al. 2016. Top-down limitation of lemmings revealed by experimental reduction of predators. - Ecology 97: 3231–3241.

Fauteux, D. et al. 2018. Evaluation of invasive and non-invasive methods to monitor rodent abundance in the Arctic. - Ecosphere 9: e02124.

Gilg, O. 2002. The summer decline of the collared lemming, Dicrostonyx groenlandicus, in high arctic Greenland. - Oikos 99: 499–510.

Gilg, O. et al. 2003. Cyclic dynamics in a simple vertebrate predator-prey community. - Science 302: 866–868.

Gilg, O. et al. 2006. Functional and numerical responses of four lemming predators in high arctic Greenland. - Oikos 113: 193–216.

Gilg, O. et al. 2009. Climate change and cyclic predator-prey population dynamics in the high Arctic. - Glob. Change Biol. 15: 2634–2652.

Gilg, O. et al. 2012. Climate change and the ecology and evolution of Arctic vertebrates. - In: Ostfeld, R. S. and Schlesinger, W. H. (eds), Year in Ecology and Conservation Biology. Blackwell Science Publ, pp. 166–190.

Gilg, O. et al. 2019. Are gastrointestinal parasites associated with the cyclic population dynamics of their arctic lemming hosts? - Int. J. Parasitol.-Parasites Wildl. 10: 6–12.

Gruyer, N. et al. 2008. Cyclic dynamics of sympatric lemming populations on Bylot Island, Nunavut, Canada. - Can. J. Zool.-Rev. Can. Zool. 86: 910–917.

Hall, L. S. et al. 1997. The Habitat Concept and a Plea for Standard Terminology. - Wildl. Soc. Bull. 1973-2006 25: 173–182.

Hansen, B. U. et al. 2008. Present-Day Climate at Zackenberg. - In: Advances in Ecological Research. High-Arctic Ecosystem Dynamics in a Changing Climate. Academic Press, pp. 111–149.

Hartig, F. 2017. DHARMa: Residual Diagnostics for Hierarchical (Multi-Level / Mixed) Regression Models. R package version 0.1.5. https://CRAN.R-project.org/package=DHARMa.

Ims, R. A. and Fuglei, E. 2005. Trophic interaction cycles in tundra ecosystems and the impact of climate change. - Bioscience 55: 311–322.

Ims, R. A. et al. 2008. Collapsing population cycles. - Trends Ecol. Evol. 23: 79–86.

Ims, R. A. et al. 2011. Determinants of lemming outbreaks. - Proc. Natl. Acad. Sci. U. S. A. 108: 1970–1974.

Kausrud, K. L. et al. 2008. Linking climate change to lemming cycles. - Nature 456: 93–U3.

Klein, D. and Bay, C. 1994. Resource partitioning by mammalian herbivores in the high arctic. - Oecologia 97: 439–450.

Mckinnon, L. et al. 2013. Predator-mediated interactions between preferred, alternative and incidental prey in the arctic tundra. - Oikos 122: 1042–1048.

Morris, D. W. et al. 2000. Measuring the ghost of competition: Insights from density-dependent habitat selection on the co-existence and dynamics of lemmings. - Evol. Ecol. Res. 2: 41–67.

Morris, D. W. et al. 2011. Forecasting ecological and evolutionary strategies to global change: an example from habitat selection by lemmings. - Glob. Change Biol. 17: 1266–1276.

Pavlis, N. K. et al. 2012. The development and evaluation of the Earth Gravitational Model 2008 (EGM2008). - J. Geophys. Res. Solid Earth 117: 1–38.

Pedersen, S. H. et al. 2016. Spatiotemporal Characteristics of Seasonal Snow Cover in Northeast Greenland from in Situ Observations. - Arct. Antarct. Alp. Res. 48: 653–671.

Porter, C. et al. 2018. ArcticDEM Release 5. Tile: 23_47_2_1_5m_v2.0_reg_dem.

Portner, H.-O. et al. 2019. IPCC Special Report on the Ocean and Cryosphere in a Changing Climate. - IPCC.

Post, E. et al. 2009. Ecological Dynamics Across the Arctic Associated with Recent Climate Change. - Science 325: 1355–1358.

Predavec, M. and Krebs, C. J. 2000. Microhabitat utilisation, home ranges, and movement patterns of the collared lemming (Dicrostonyx groenlandicus) in the central Canadian Arctic. - Can. J. Zool.-Rev. Can. Zool. 78: 1885–1890.

Prins, H. H. Th. 1982. Why Are Mosses Eaten in Cold Environments Only? - Oikos 38: 374–380.

Prost, S. et al. 2010. Influence of Climate Warming on Arctic Mammals? New Insights from Ancient DNA Studies of the Collared Lemming Dicrostonyx torquatus. - Plos One 5: e10447.

Prost, S. et al. 2013. Losing ground: past history and future fate of Arctic small mammals in a changing climate. - Glob. Change Biol. 19: 1854–1864.

R Core Team 2020. R: A Language and Environment for Statistical Computing. - R Foundation for Statistical Computing.

Rau, F. 1995. Geoökologische und hydrologische Untersuchungen in einem hochaktischen Tundrenökosystem auf Traill Ø, Nordost-Grönland. Unpublished diploma thesis. Albert Ludwig Universität Freiburg, Germany.

Reid, D. G. and Krebs, C. J. 1996. Limitations to collared lemming population growth in winter. - Can. J. Zool.-Rev. Can. Zool. 74: 1284–1291.

Reid, D. G. et al. 2012. Lemming winter habitat choice: a snow-fencing experiment. - Oecologia 168: 935–946.

Rodgers, A. and Lewis, M. 1985. Diet selection in arctic lemmings (lemmus-Sibericus and Dicrostonyx-Groenlandicus) - food preferences. - Can. J. Zool.-Rev. Can. Zool. 63: 1161–1173.

Rodgers, A. and Lewis, M. 1986. Diet selection in arctic lemmings (lemmus-Sibiricus and Dicrostonyx-Groenlandicus) - demography, home range, and habitat use. - Can. J. Zool.-Rev. Can. Zool. 64: 2717–2727.

Schmidt, N. M. 2000. Spatiotemporal distribution and habitat use of the collared lemming, Dicrostonyx groenlandicus Traill, in high arctic Northeast Greenland.

Schmidt, N. et al. 2012a. High Arctic plant community responses to a decade of ambient warming. - Biodiversity 13: 191–199.

Schmidt, N. M. et al. 2012b. Response of an arctic predator guild to collapsing lemming cycles. - Proc. R. Soc. B-Biol. Sci. 279: 4417–4422.

Schmidt, N. M. et al. 2019. An ecosystem-wide reproductive failure with more snow in the Arctic. - PLOS Biol. 17: e3000392.

Sittler, B. 1995. Response of stoats (mustela-Erminea) to a fluctuating lemming (dicrostonyx-Groenlandicus) population in North-East Greenland - Preliminary results from a long-term study. - Ann. Zool. Fenn. 32: 79–92.

Sittler, B. et al. 2000. Low abundance of king elder nests during low lemming years in northeast Greenland. - Arctic 53: 53–60.

Soininen, E. M. et al. 2015. Highly overlapping winter diet in two sympatric lemming species revealed by DNA metabarcoding. - Plos One 10: e0115335.

Therrien, J. F. et al. 2014. Predation pressure by avian predators suggests summer limitation of small-mammal populations in the Canadian Arctic. - Ecology 95: 56–67.

Turchin, P. et al. 2000. Are lemmings prey or predators? - Nature 405: 562–565.

Walker, D. A. et al. 2005. The Circumpolar Arctic vegetation map. - J. Veg. Sci. 16: 267–282.

Wookey, P. A. et al. 2009. Ecosystem feedbacks and cascade processes: understanding their role in the responses of Arctic and alpine ecosystems to environmental change. - Glob. Change Biol. 15:1153–1172.

