## Supporting information for "Long-term monitoring reveals topographical features and vegetation explain winter habitat use of an Arctic rodent"

### Data cleaning

Data cleaning was necessary as data on the position of lemming nests has not been used yet. Data cleaning involved fixing obvious typing errors, such as transposed digits, missing values, and deleting empty rows. The preliminary data management and cleaning was done with Microsoft Excel 2016.

### Vegetation digitization

Vegetation data was derived from paper maps, which had to be scanned and georeferenced. Next, a digital version was created by cutting the vegetation zones into polygons (Fig. A6). Vegetation data for each patch was calculated by splitting the polygon area by the grid and assigning it to the patches (see Supplementary material Appendix 1, Fig. A6). For each patch, we then calculated the coverage of the six vegetation types.

### Method discussion & further research needs

In contrast to other studies mapping and analyzing lemming winter nests, our approach differs due to the origin of the underlying data, which comes with a certain amount of inaccuracy. Instead of assessing habitat data like slope, elevation and vegetation in the field (Duchesne et al., 2011), we used a digital elevation model and a vegetation map to analyze habitat selection. Furthermore, winter nests were not located by GPS-coordinates (Reid et al., 2012), but instead allocated to single land patches of 6.25ha in size (see Methods). These patches can contain myriad topographic elements (Fig. A11, Fig. A12, Fig. A13) and thus a mean value may not be able to represent the whole area correctly. Additionally, the exact locations of nests inside a certain patch are unknown, which means that nests may be truly associated with a vegetation/slope minority and thus are not represented in the statistical analysis. Moreover, the process of digitization may have shifted nest positions slightly. While the georeferencing process was accurate, the digital grid did not align perfectly on all aerial photos (Appendix). Still, remote sensing and geographic information systems offer great potential for future lemming winter ecology assessment. Further research on Traill Island should make the transition to record winter nests with exact GPS-positions and create digital buffers around individual nests to analyze habitat features in close proximity.

Micro-topography is an important factor in determining winter habitat selection because it increases the probability of subnivean air space formation by affecting the pattern of snow drift at a small scale and provides a refuge against subnivean flooding (Duchesne et al., 2011). Analyzing hummocky areas probably requires a more accurate digital elevation model than that available for this study. Drones with mounted cameras are a cost-efficient method in acquiring elevation models with superior resolutions (Bernard et al., 2017). Furthermore, it is suggested to analyze geomorphological structures, such as terraces and gullies, as lemmings seem to favour them for their winter habitats. Terraces are features that can be automatically identified with the TerEx Terrace Extraction tool (Stout and Belmont, 2014).

Precise snow depth data has an enormous potential for further research as it is the main factor influencing winter nest distribution (Reid and Krebs, 1996). Previous lemming studies either measured snow depth at selected points with a metal rod (Duchesne et al., 2011; Reid et al., 2012) or modelled the snow conditions with the SNOWPACK software using meteorological data as input variables (Bilodeau et al., 2013). However, since the spatial distribution of snow in the Arctic determine the landscape patterns, the latter method is unable to model local distribution. We therefore suggest to apply a different model which also accounts for blowing-snow redistribution and sublimation, such as Alpine3D (Lehning et al., 2006). In any case, access to local weather data is crucial for future snow modelling ambitions and would require installing a weather station in the study area.

Further research may also involve analyzing vegetation cover in proximity to individual winter nests through remote sensing. Previous studies noted that satellite imagery has great potential as a tool for quantifying and monitoring biophysical variables in the High Arctic. The vegetation index, assessed with multispectral satellite bands, was highly correlated with percent cover collected in the field on Boothia Peninsula, Canada (Laidler et al., 2008). Modern satellite imagery is able to differentiate between vegetation types and even moisture content in the Arctic (Liu et al., 2017), with a resolution that allows identifying habitat features of small mammals.

Further research should also evaluate habitat selection by combining data on nest sites and vegetation availability, for instance by using the ratio of mean proportion of the vegetation in proximity of the nest over the mean availability (Soininen et al., 2015). Additionally, the effects shown in this study may not be linear. Lemmings may prefer middle values of some predictors, for example slope and terrain ruggedness, as they would represent different topographic elements. This could be tested by including quadratic terms in the statistical analysis.

| 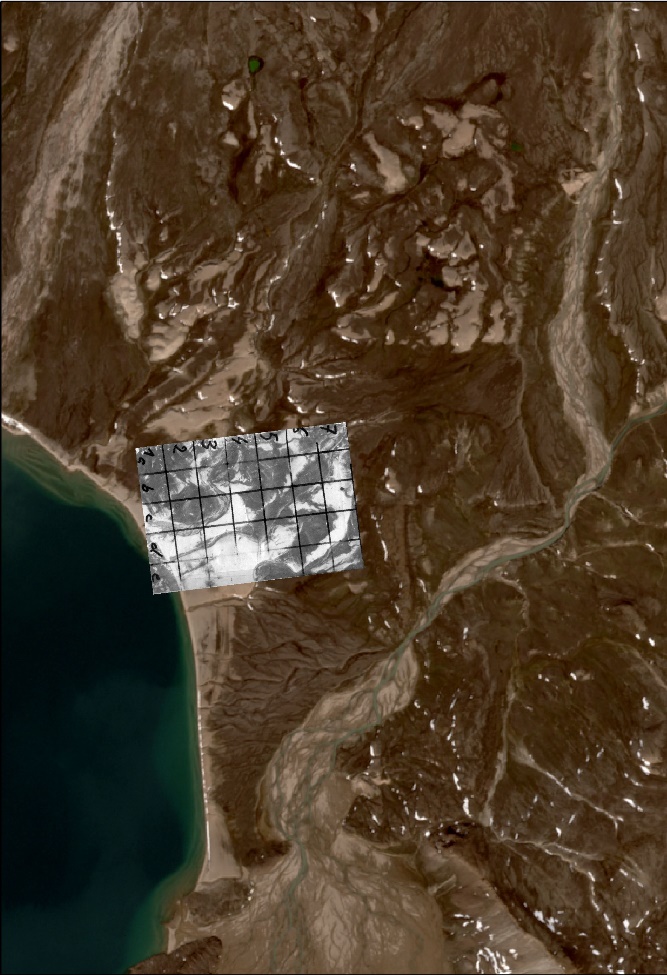  Figure 1. Georeferenced aerial photo (one of eleven). | 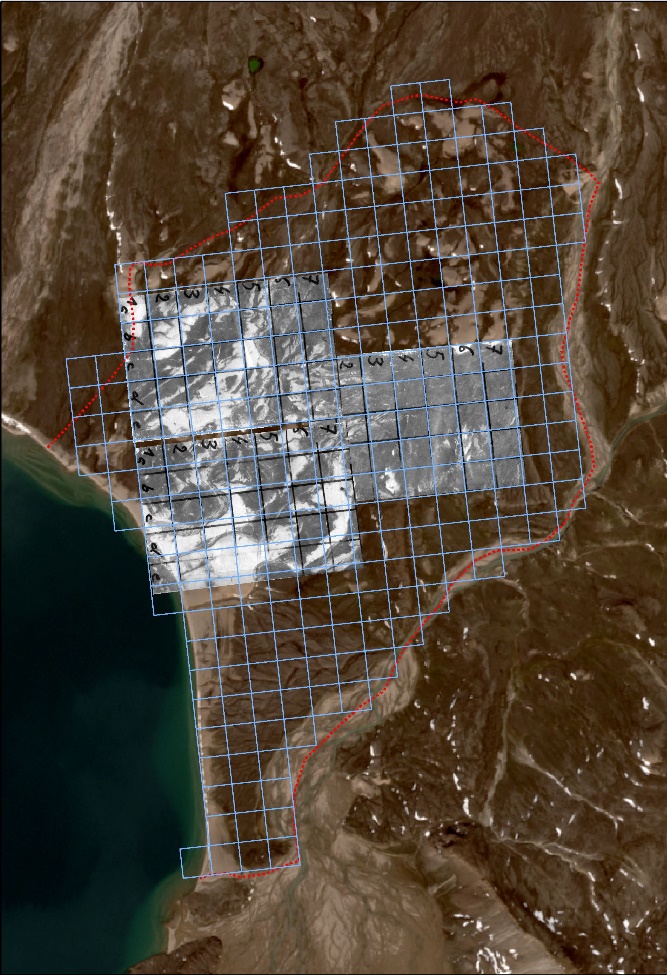  Figure 2. Final digital grid only including patches with data and study area borders. The three sheets overlap in the centre of the image. |
| --- | --- |
| 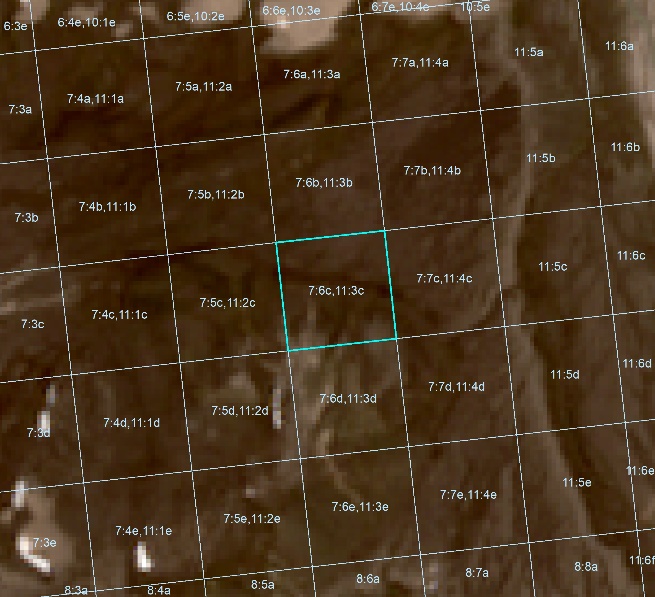  Figure 3. Old patch identification system based on aerial photo sheet numbers and rows/column of single sheets. Patch 7:6c, 11:3c comprises two overlapping aerial photos. | 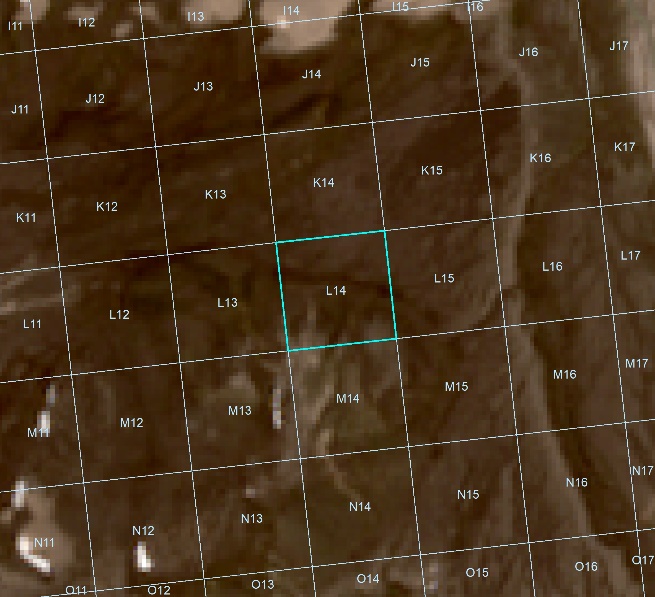  Figure 4. New patch identification system includes merged aerial photos. Patch 7:6c, 11:3c was transformed to L14 and the data combined. In total, 212 winter nests were found on this patch. |

| 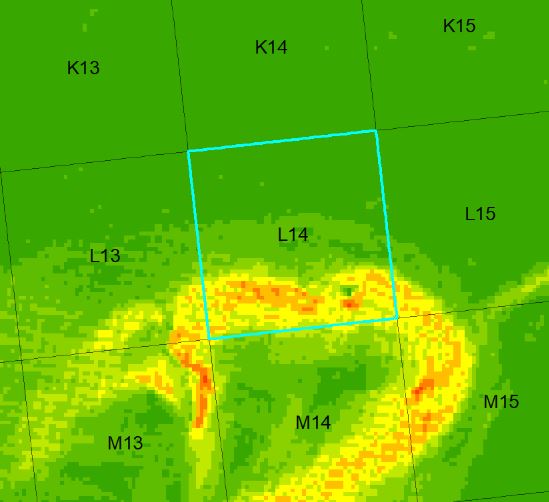  Figure 5. Slope values of patch L14; min:0,3% max:27,8% mean:5,2%. | 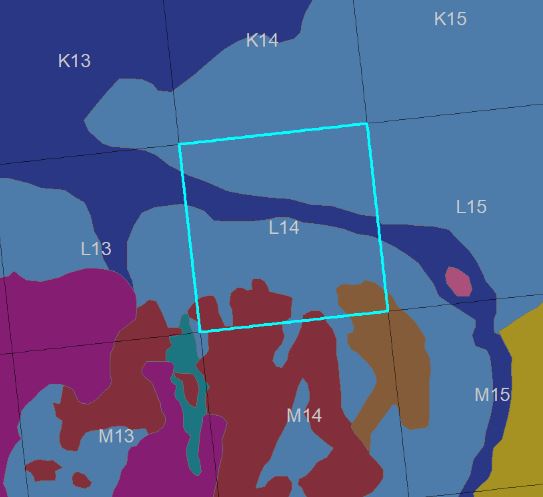  Figure 6. Vegetation data for patch L14, 47,068m² (75%) spot tundra, 7,376m² (12%) moss-sedges tundra, 2458m² (4%) solifluidal vegetation-free zones and 5,620m² (9%) clay bumps. |
| --- | --- |

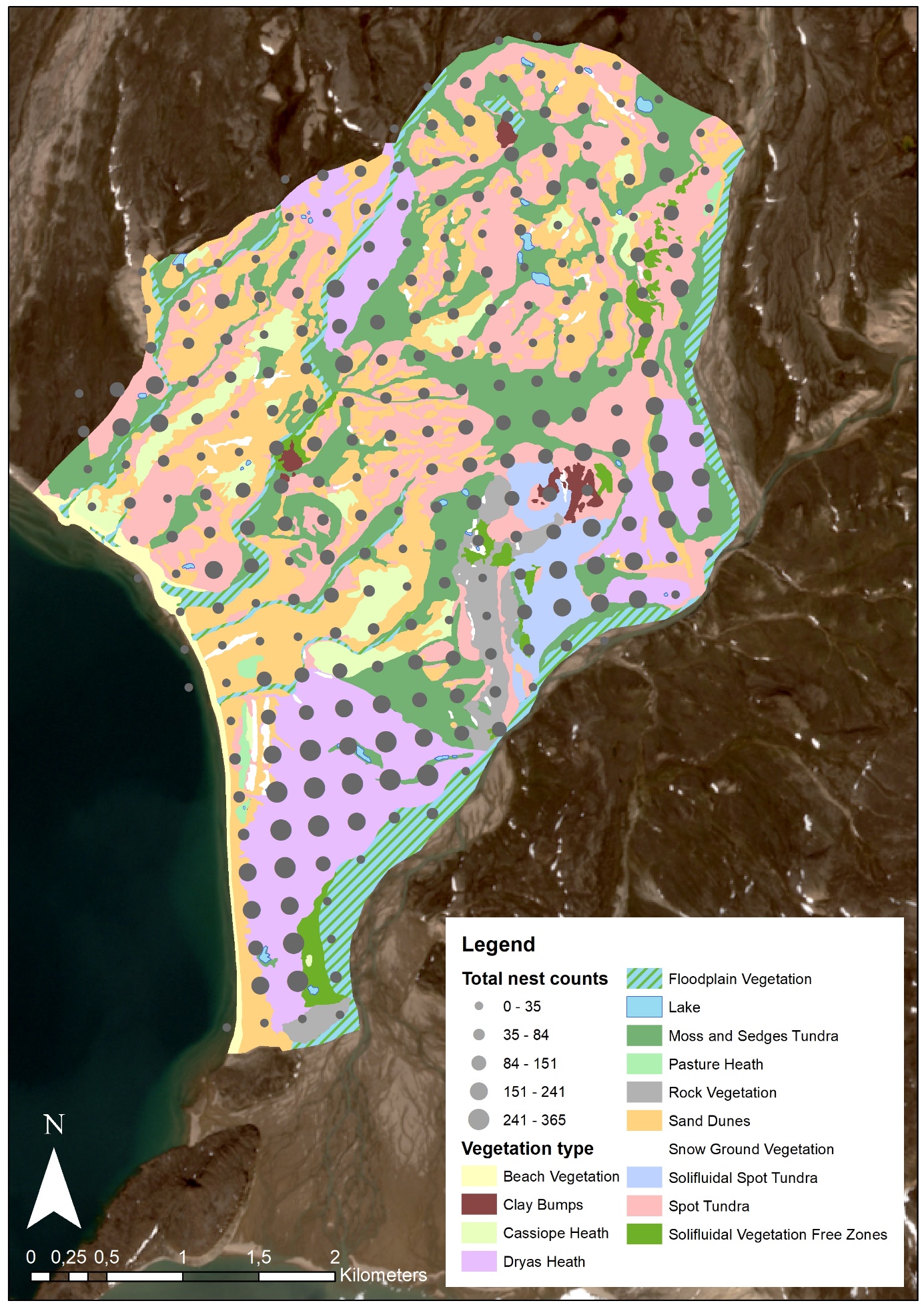

Figure 7. Vegetation data of the study area.

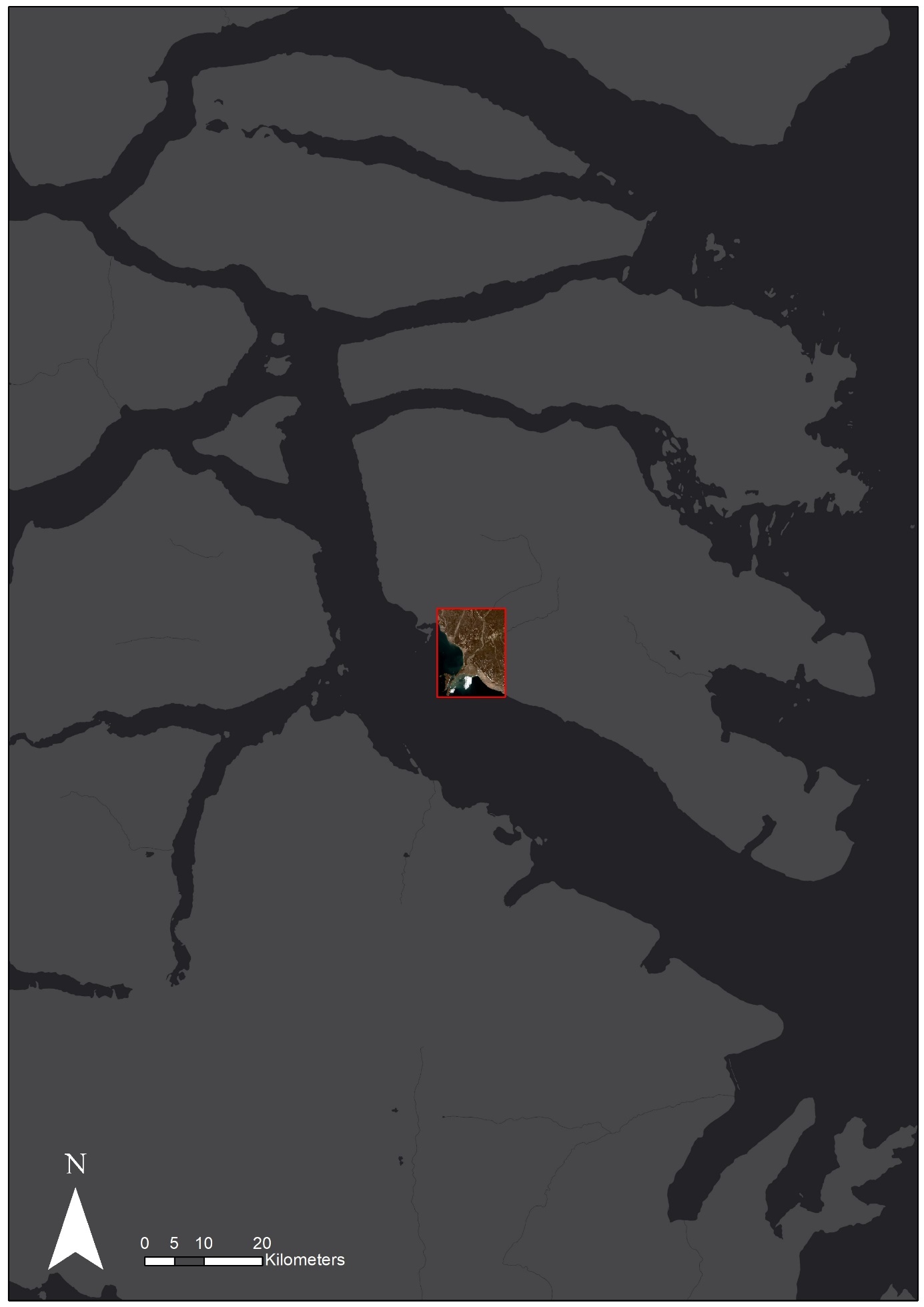

Figure 8. Location of the study area on Traill island, Greenland.

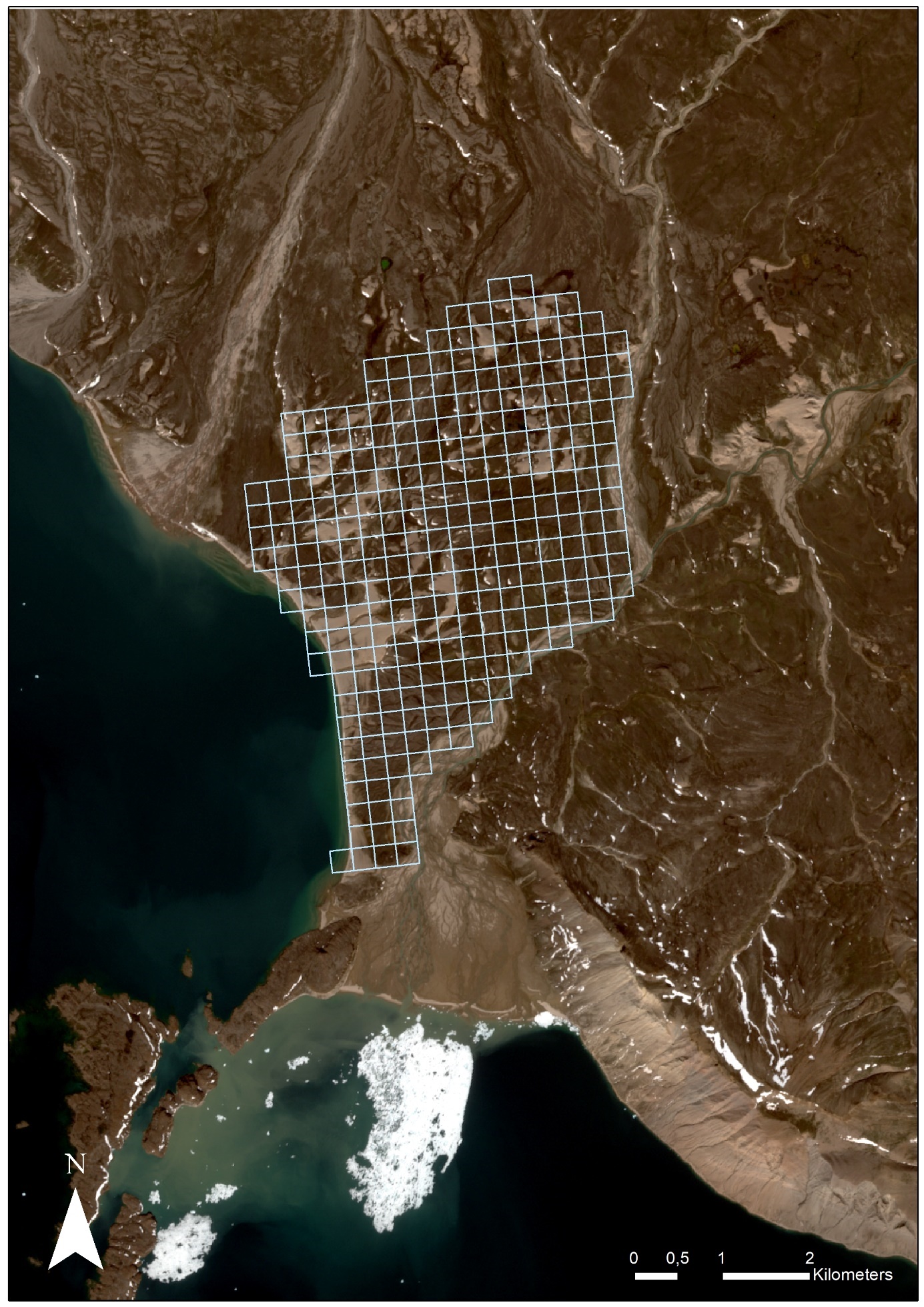

Figure 9. Study grid.

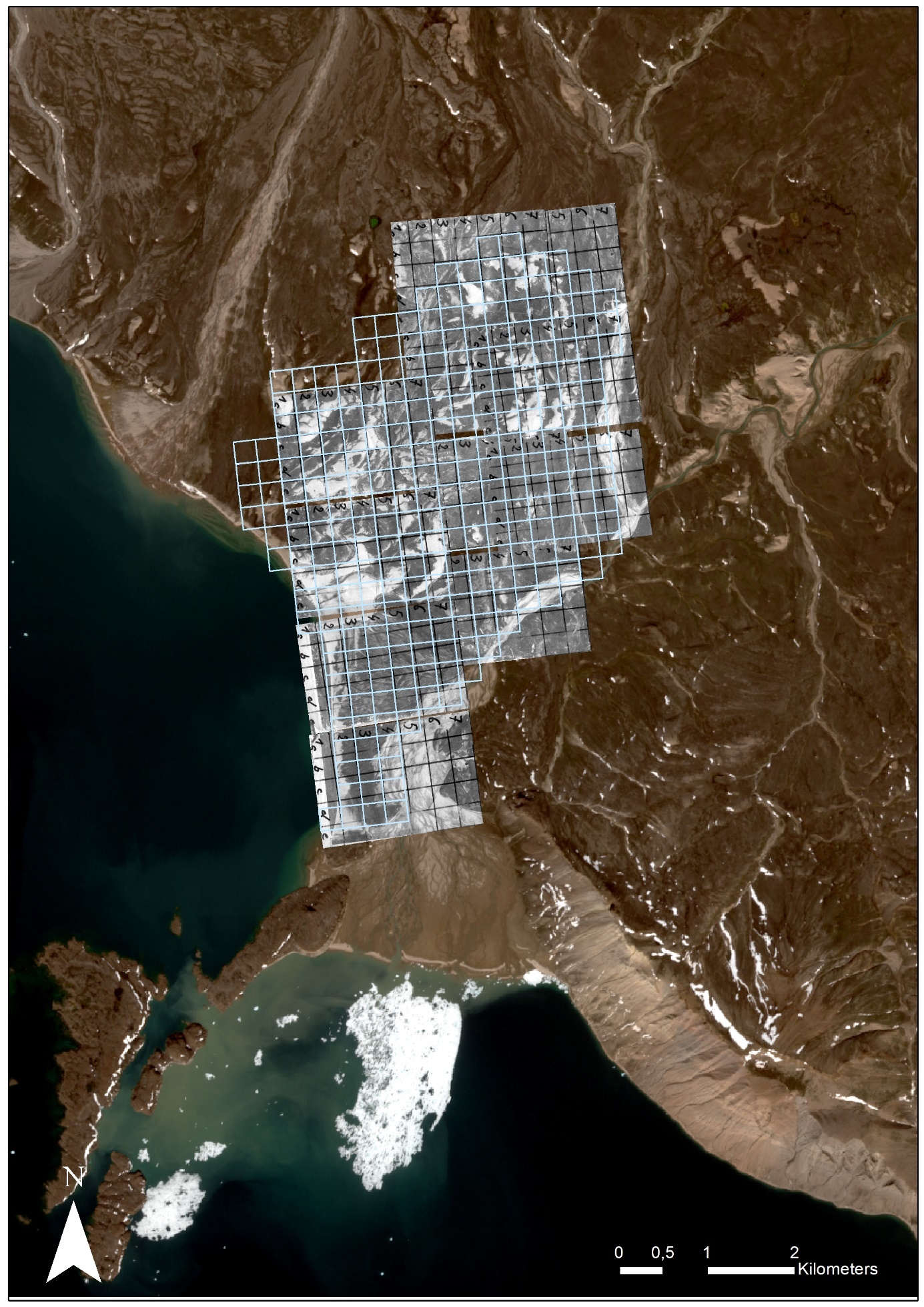

Figure 10. Study grid derived from aerial photos.

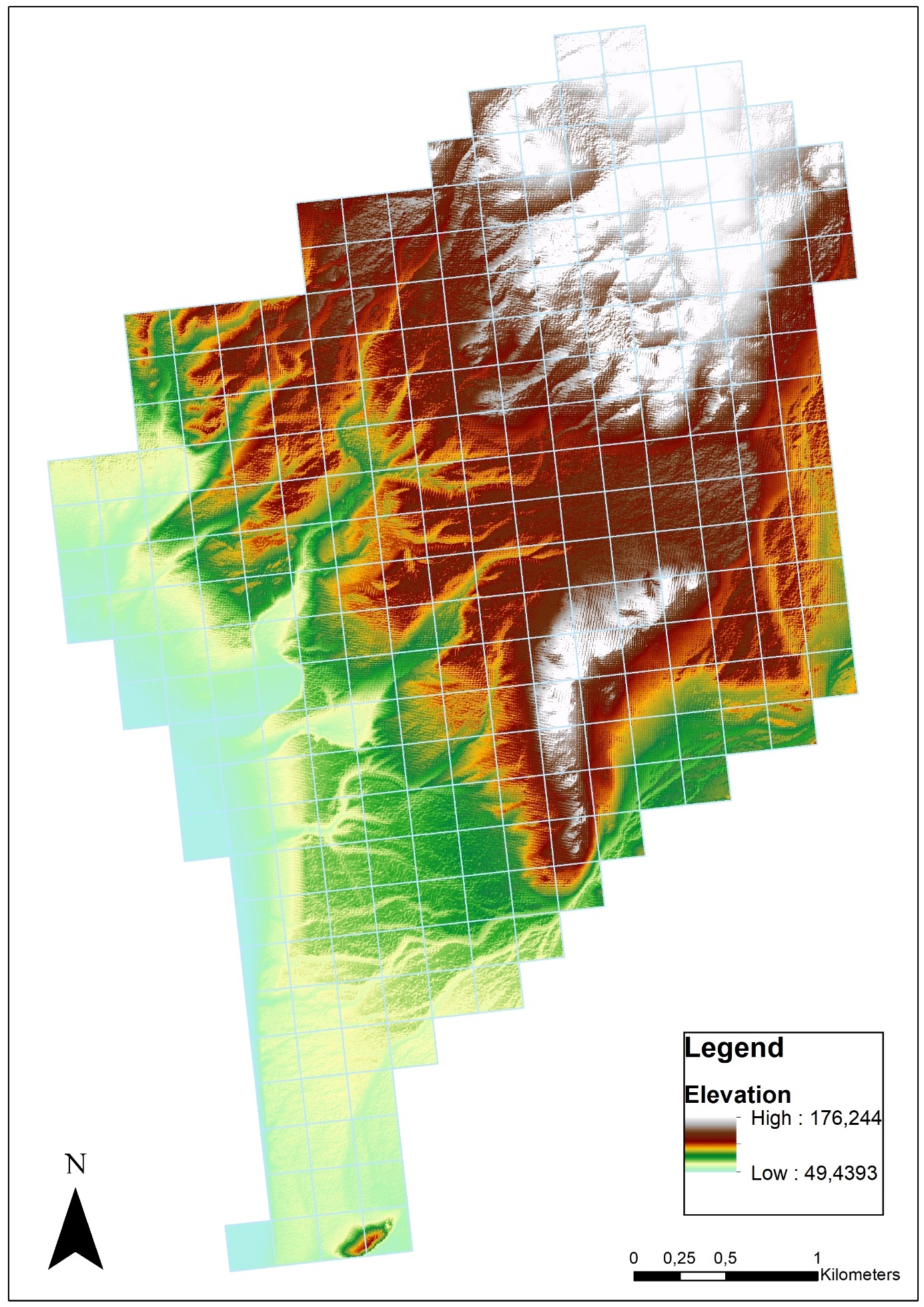

Figure 11. Elevation of the study area.

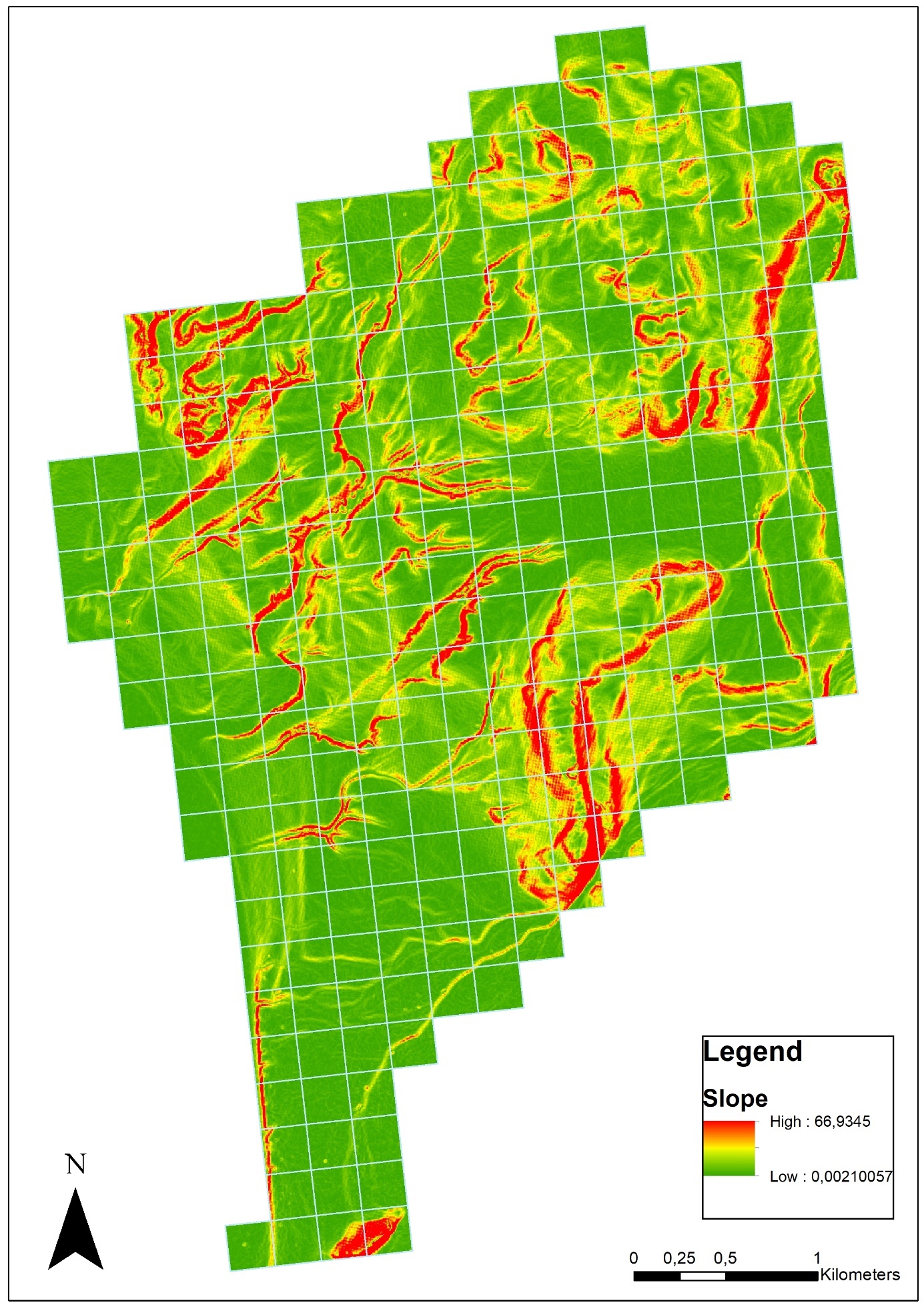

Figure 12. Slope of the study area.

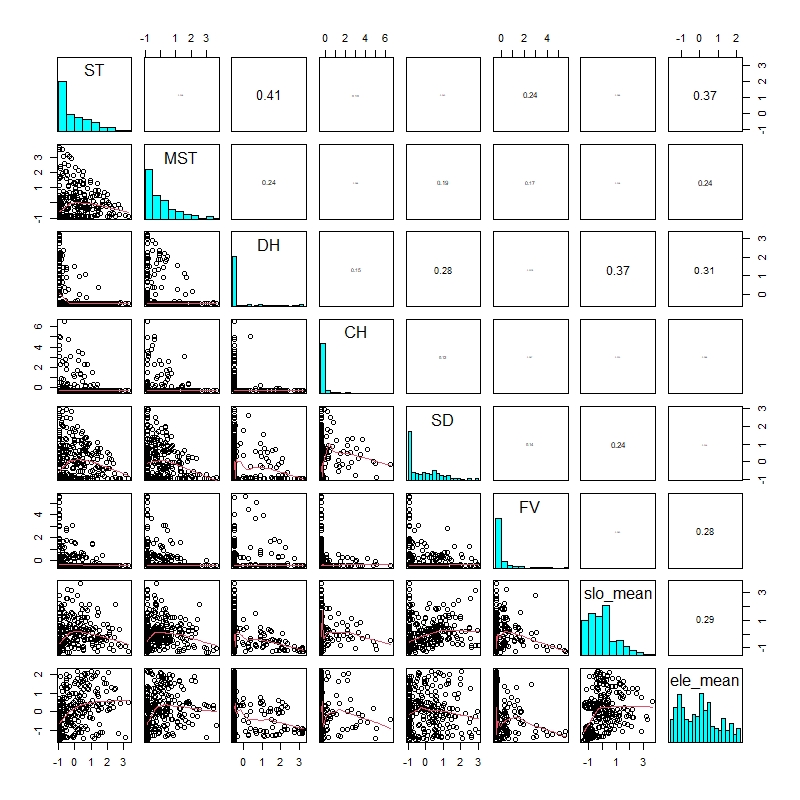

Figure 13. Pairs plot of the predictors. All correlation coefficients are smaller than 0.41.

Table 1. Summary table of negative binomial generalized linear model of the effect of vegetation type, elevation and slope on total lemming nest number in low density (< 2 lemmings/ha) years. Significant results are highlighted in bold.

| Fixed Factors | Estimate | Std. Error | Z-value | P-value |
| --- | --- | --- | --- | --- |
| Intercept | **-1.21709** | **0.20355** | **-5.979** | **<0.001** |
| Spot Tundra | **0.28119** | **0.08556** | **3.287** | **0.001** |
| Moss-Sedges Tundra | **0.23675** | **0.07843** | **3.018** | **0.003** |
| Dryas Heath | **0.62336** | **0.09434** | **6.607** | **<0.001** |
| Cassiope Heath | -0.03010 | 0.07241 | -0.416 | 0.678 |
| Sand Dunes | -0.14770 | 0.08026 | -1.840 | 0.0657 |
| Floodplain vegetation | **-0.15366** | **0.07732** | **-1.987** | **0.047** |
| Mean Slope | **0.26044** | **0.07678** | **3.392** | **<0.001** |
| Mean Elevation | **-0.44014** | **0.09636** | **-4.568** | **<0.001** |

Table 2. Summary table of negative binomial generalized linear model of the effect of vegetation type, elevation and slope on total lemming nest number in high density (>2 lemmings/ha) years. Significant results are highlighted in bold.

| Fixed Factors | Estimate | Std. Error | Z-value | P-value |
| --- | --- | --- | --- | --- |
| Intercept | **0.79899** | **0.20340** | **3.928** | **<0.001** |
| Spot Tundra | **0.38152** | **0.06786** | **5.623** | **<0.001** |
| Moss-Sedges Tundra | **0.33627** | **0.06227** | **5.400** | **<0.001** |
| Dryas Heath | **0.71427** | **0.07676** | **9.305** | **<0.001** |
| Cassiope Heath | 0.07254 | 0.05495 | 1.320 | 0.187 |
| Sand Dunes | -0.02137 | 0.06429 | -0.332 | 0.739 |
| Floodplain vegetation | **-0.18456** | **0.06303** | **-2.928** | **0.003** |
| Mean Slope | **0.23401** | **0.06061** | **3.861** | **<0.001** |
| Mean Elevation | **-0.29629** | **0.09229** | **-3.210** | **0.001** |

Table 3. Vegetation description of the lower Karupelv-Valley, Traill O. (from Rau 1995, translated and modified).

| **Vegetation Type** | **Total Surface Area in m^2^ (%)** | **Description** | **Surface Coverage** |
| --- | --- | --- | --- |
| Cassiope heath | 654097 (4,04) | Dwarf bushes with pure stands of Cassiope tetragona, as well as numerous other dwarf shrubs (e.g. *Salix arctica*, *Betula nana*). Climax vegetation of the northeast Greenland coastal area. | up to 100% |
| Dryas heath | 2334351 (14,45) | Dwarf shrub heaths are dominated by *Dryas octopetala*, along with numerous upholstery and rosette plants. | between 31-60% |
| Spot tundra | 3726263 (22,80) | Due to sparse vegetation with sporadically occurring vegetation islands of Cassiope heath and Dryas heath embossed vegetation unit. The interspaces are occupied by moss-lichen societies. | between 31-60% |
| Solifluidal spot tundra | 433274 (2,68) | Variation of the spot tundra on solifluidally oriented locations (solifluction lugs, garland bottoms, etc.). | between 31-60% |
| Pasture heath | 64008 (0,40) | Vegetation characterized by steppe with predominant grasses (*Poaceae*) and sour grasses (e.g. *Carex spec*., *Kobresia myosuroidas*), as well as *Dryas octopetala* and *Salix actica*. | between 31-60% |
| Moss and sedges tundra | 3410119 (20,36) | Wet vegetation dominated by moose and sedges (Carex spec.), often with cottongrass (*Eriopherum spec.*). Species-rich and mostly high-growing communities, dwarf shrubs are largely missing. | between 31-90% |
| Floodplain vegetation | 1204318 (5,74) | Species- and individual poor recent floodplains. Characteristic are various species of rockfoils (*Saxifraga aizoides, S. nathorstii*), on fine material also cottongrass (*Eriophorum scheuchzeri*) and rushes (*Juncus spec.*). | less than 10% |
| Beach vegetation | 214800 (0,99) | Species- and individual poor, halophilic vegetation with seawater influence. Isolated upholstery and rosette plants (e.g., *Cochlearia groenlandica, Honkenya peploides*). | less than 10% |
| Snow ground vegetation | 157216 (0,98) | For the most part, small-scale populations of long-lasting snow cover sites, moss hollows with isolated vascular plants (e.g., *Oxyria digyna, Ranunculus pygmaeus*). | less than 30% |
| Rock vegetation | 443041 (2,68) | Vegetation unit bound to adjacent rocks and block debris with numerous polyporous plants (such as *Silene acaulis*) and isolated Cassiope mats. In damp crevasses also *Sedum roseum*, more rarely *Cystopteris fragilis* and *Woodsia glabella*. |  |
| Sand dunes | 3597133 (21,94) | Individually poor, largely vegetation-free stocks (polar semi-deserts). Isolated upholstery and rosette plants (e.g., *Silene acaulis*, *Papaver radicatum*) often have a low-grade crust consisting of mosses and lichens. | less than 10% |
| Clay-bumps | 161,30 (0,62) | Vegetation-free stands on clay bumps, in dry cracks trellis-growth of *salix actica*. | less than 10% |
| Solifluidal vegetation free zones | 310702 (1,93) | Completely vegetation-free areas with strong, mostly amorphous solifluction. Isolated upholstery plants at favorable sites. | less than 5% |

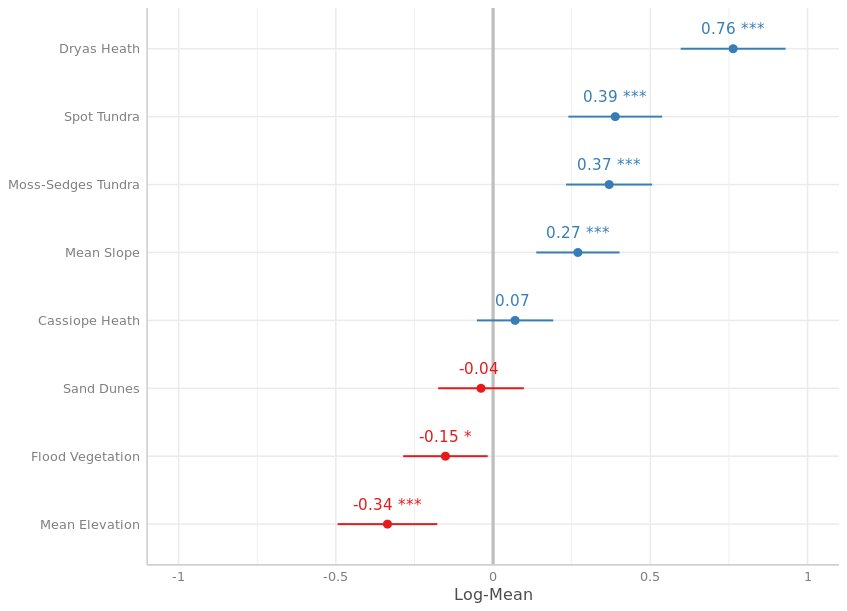

Figure 14: Forest plot for all significant predictors and for all years between 1989 and 2019.

Table 4. Empty patches.

| **Year** | **Total Nests** | **Occupied grids** |
| --- | --- | --- |
| 1989 | 1285 | 151 |
| 1990 | 3685 | 213 |
| 1991 | 286 | 109 |
| 1992 | 103 | 41 |
| 1993 | 144 | 58 |
| 1994 | 2411 | 202 |
| 1995 | 1007 | 177 |
| 1996 | 195 | 77 |
| 1997 | 311 | 81 |
| 1998 | 3469 | 208 |
| 1999 | 1598 | 195 |
| 2000 | 111 | 59 |
| 2001 | 302 | 96 |
| 2002 | 403 | 119 |
| 2003 | 59 | 39 |
| 2004 | 889 | 130 |
| 2005 | 211 | 94 |
| 2006 | 778 | 119 |
| 2007 | 221 | 81 |
| 2008 | 272 | 57 |
| 2009 | 21 | 13 |
| 2010 | 290 | 75 |
| 2011 | 1067 | 146 |
| 2012 | 1338 | 160 |
| 2013 | 69 | 48 |
| 2014 | 30 | 21 |
| 2015 | 271 | 84 |
| 2016 | 457 | 124 |
| 2017 | 987 | 142 |
| 2018 | 44 | 20 |
| 2019 | 455 | 110 |

Table 5. Moran I test for spatial autocorrelation with observed, expected, standard errors and P-values. Significant (P≤0.05) years are printed in bold.

| Year | Observed | Expected | Std. Error | P |
| --- | --- | --- | --- | --- |
| 1989 | **0.0336845** | **-0.003663004** | **0.004742371** | **3.330669e-15** |
| 1990 | **0.03711665** | **-0.003663004** | **0.004726747** | **0** |
| 1991 | **0.008084821** | **-0.003663004** | **0.004741589** | **0.01322642** |
| 1992 | -0.006072568 | -0.003663004 | 0.004742053 | 0.6113645 |
| 1993 | **0.01057907** | **-0.003663004** | **0.004740968** | **0.002664164** |
| 1994 | **0.03253545** | **-0.003663004** | **0.004732132** | **2.020606e-14** |
| 1995 | **0.01620116** | **-0.003663004** | **0.00473879** | **2.767223e-05** |
| 1996 | -0.002687771 | -0.003663004 | 0.004743358 | 0.8371037 |
| 1997 | **0.02233693** | **-0.003663004** | **0.004744456** | **4.251684e-08** |
| 1998 | **0.03339672** | **-0.003663004** | **0.004725563** | **4.440892e-15** |
| 1999 | **0.009420346** | **-0.003663004** | **0.004734185** | **0.005716916** |
| 2000 | **0.007260295** | **-0.003663004** | **0.004743228** | **0.02128306** |
| 2001 | **0.0264147** | **-0.003663004** | **0.004743132** | **2.278078e-10** |
| 2002 | **0.00755862** | **-0.003663004** | **0.00474194** | **0.017959** |
| 2003 | -0.001897081 | -0.003663004 | 0.004739422 | 0.709444 |
| 2004 | **0.02397695** | **-0.003663004** | **0.004744107** | **5.671544e-09** |
| 2005 | -0.001597497 | -0.003663004 | 0.004744279 | 0.6632954 |
| 2006 | 0.003710582 | -0.003663004 | 0.004743095 | 0.1200429 |
| 2007 | **0.006487589** | **-0.003663004** | **0.004744159** | **0.03238727** |
| 2008 | **0.02997269** | **-0.003663004** | **0.004742426** | **1.316947e-12** |
| 2009 | **0.008166707** | **-0.003663004** | **0.004740444** | **0.01257849** |
| 2010 | **0.01369994** | **-0.003663004** | **0.004743238** | **0.0002516562** |
| 2011 | **0.01352658** | **-0.003663004** | **0.004743301** | **0.0002901133** |
| 2012 | **0.01166813** | **-0.003663004** | **0.004740957** | **0.001221709** |
| 2013 | **0.01025571** | **-0.003663004** | **0.004742402** | **0.003336004** |
| 2014 | -0.0008750164 | -0.003663004 | 0.004737033 | 0.5561623 |
| 2015 | 0.005210595 | -0.003663004 | 0.004742303 | 0.06132318 |
| 2016 | **0.006076428** | **-0.003663004** | **0.004743759** | **0.04006247** |
| 2017 | **0.02065061** | **-0.003663004** | **0.004744543** | **2.982624e-07** |
| 2018 | -0.002603623 | -0.003663004 | 0.004739246 | 0.8231202 |
| 2019 | **0.019045** | **-0.003663004** | **0.004744086** | **1.696372e-06** |

### References

Bernard, É., Friedt, J.M., Tolle, F., Griselin, M., Marlin, C., Prokop, A., 2017. Investigating snowpack volumes and icing dynamics in the moraine of an Arctic catchment using UAV photogrammetry. Photogramm. Rec. 32, 497–512. https://doi.org/10.1111/phor.12217

Bilodeau, F., Gauthier, G., Berteaux, D., 2013. The effect of snow cover on lemming population cycles in the Canadian High Arctic. Oecologia 172, 1007–1016. https://doi.org/10.1007/s00442-012-2549-8

Duchesne, D., Gauthier, G., Berteaux, D., 2011. Habitat selection, reproduction and predation of wintering lemmings in the Arctic. Oecologia 167, 967–980. https://doi.org/10.1007/s00442-011-2045-6

Laidler, G.J., Treitz, P.M., Atkinson, D.M., 2008. Remote sensing of arctic vegetation: Relations between the NDVI, spatial resolution and vegetation cover on Boothia Peninsula, Nunavut. Arctic 61, 1–13. https://doi.org/10.14430/arctic2

Lehning, M., Völksch, I., Gustafsson, D., Nguyen, T.A., Stähli, M., Zappa, M., 2006. ALPINE3D: a detailed model of mountain surface processes and its application to snow hydrology. Hydrol. Process. 20, 2111–2128. https://doi.org/10.1002/hyp.6204

Liu, N., Budkewitsch, P., Treitz, P., 2017. Examining spectral reflectance features related to Arctic percent vegetation cover: Implications for hyperspectral remote sensing of Arctic tundra. Remote Sens. Environ. 192, 58–72. https://doi.org/10.1016/j.rse.2017.02.002

Reid, D.G., Bilodeau, F., Krebs, C.J., Gauthier, G., Kenney, A.J., Gilbert, B.S., Leung, M.C.-Y., Duchesne, D., Hofer, E., 2012. Lemming winter habitat choice: a snow-fencing experiment. Oecologia 168, 935–946. https://doi.org/10.1007/s00442-011-2167-x

Reid, D.G., Krebs, C.J., 1996. Limitations to collared lemming population growth in winter. Can. J. Zool.-Rev. Can. Zool. 74, 1284–1291. https://doi.org/10.1139/z96-143

Soininen, E.M., Gauthier, G., Bilodeau, F., Berteaux, D., Gielly, L., Taberlet, P., Gussarova, G., Bellemain, E., Hassel, K., Stenoien, H.K., Epp, L., Schroder-Nielsen, A., Brochmann, C., Yoccoz, N.G., 2015. Highly overlapping winter diet in two sympatric lemming species revealed by DNA metabarcoding. Plos One 10, e0115335. https://doi.org/10.1371/journal.pone.0115335

Stout, J.C., Belmont, P., 2014. TerEx Toolbox for semi-automated selection of fluvial terrace and floodplain features from lidar. Earth Surf. Process. Landf. 39, 569–580. https://doi.org/10.1002/esp.3464
